# 3D Epigenomic Characterization Reveals Insights Into Gene Regulation and Lineage Specification During Corticogenesis

**DOI:** 10.1101/2020.02.24.963652

**Authors:** Michael Song, Mark-Phillip Pebworth, Xiaoyu Yang, Armen Abnousi, Changxu Fan, Jia Wen, Jonathan D. Rosen, Mayank NK Choudhary, Xiekui Cui, Ian R. Jones, Seth Bergenholtz, Ugomma C. Eze, Ivan Juric, Bingkun Li, Lenka Maliskova, Weifang Liu, Alex A. Pollen, Yun Li, Ting Wang, Ming Hu, Arnold R. Kriegstein, Yin Shen

## Abstract

Lineage-specific epigenomic changes during human corticogenesis have previously remained elusive due to challenges with tissue heterogeneity and sample availability. Here, we analyze cis-regulatory chromatin interactions, open chromatin regions, and transcriptomes for radial glia, intermediate progenitor cells, excitatory neurons, and interneurons isolated from mid-gestational human brain samples. We show that chromatin looping underlies transcriptional regulation for lineage-specific genes, with transcription factor motifs, families of transposable elements, and disease-associated variants enriched at distal interacting regions in a cell type-specific manner. A subset of promoters exhibit unusually high degrees of chromatin interactivity, which we term super interactive promoters. Super interactive promoters are enriched for critical lineage-specific genes, suggesting that interactions at these loci contribute to the fine-tuning of cell type-specific transcription. Finally, we present CRISPRview, a novel approach for validating distal interacting regions in primary cells. Our study presents the first characterization of cell type-specific 3D epigenomic landscapes during human corticogenesis, advancing our understanding of gene regulation and lineage specification during human brain development.

## Introduction

The human cortex is a complex, heterogeneous structure that undergoes extensive expansion during development, a process which is markedly different and features distinct cell types from mouse cortical development. Previous studies utilized single cell RNA sequencing (scRNA-seq) to unravel the transcriptomic diversity of the developing cortex, revealing at least nine major cell types and up to 26 distinct subtypes in the dorsal cortex alone^1,2^. Much of this diversity arises from cortical stem cells known as radial glia (RG), whose cell bodies reside in the germinal zones (GZs) of the dorsal and ventral cortex. In the dorsal cortex, RG divide asymmetrically to give rise to intermediate progenitor cells (IPCs), which proliferate and differentiate into excitatory neurons (eNs)^3,4^. These newborn neurons undergo radial migration until they reach the cortical plate (CP), where they mature and undergo synaptogenesis^5^. Meanwhile, interneurons (iNs) produced in the ventral cortex migrate tangentially into the dorsal cortex through the marginal and germinal zones^6^. These processes result in a CP consisting primarily of eNs and iNs and a GZ where all four cell types are intermixed.

Dynamic changes in the epigenomic landscape have been shown to play a critical role in development and cell fate commitment, for instance through the rewiring of physical chromatin loops between promoters and distal regulatory elements^7^. These regulatory interactions are of particular interest as their dysregulation has been linked to complex diseases and traits^8,9^. Despite their utility, detailed epigenomic characterizations are still absent for specific cell types in the developing cortex due to shortcomings associated with the analysis of bulk tissues^10,11^. Here, we present a novel strategy for isolating RG, IPCs, eNs, and iNs from mid-gestational human brain samples, enabling the cell type-specific profiling of their epigenomic features. In addition, we present CRISPRview, a technique for validating distal regulatory regions in primary cells, demonstrating that *GPX3*, *TNC*, and *HES1* are regulated by distal enhancers in RG. Our results identify novel mechanisms underlying gene regulation and lineage specification during corticogenesis, providing a framework for the elucidation of diverse processes in development and disease.

## Results

### Isolation of specific cell populations from the developing human cortex

To isolate four specific cell populations (RG, IPCs, eNs, and iNs) from mid-gestational human brain samples between gestational weeks (GW) 15 to 22 (**Supplementary Table 1**), we adapted a previously reported approach for isolating RG from human cortical samples using fluorescence-activated cell sorting (FACS)^12^. Specifically, we incorporated markers for additional cell types from a recently published scRNA-seq dataset in the human neocortex^1^. Microdissected GZ and CP samples were dissociated, stained using antibodies for EOMES, SOX2, PAX6, and SATB2, then partitioned into their constituent populations using FACS (**Fig. 1a and Supplementary Fig. 1**). IPCs were first isolated as the EOMES+ population. eNs were isolated from the EOMES- and SOX2-population based on high expression of SATB2, which marks both newborn and mature eNs at the ages of the samples^1^. RG were isolated based on the high expression of both SOX2 and PAX6, and iNs were isolated based on medium SOX2 and low PAX6 expression.

**Figure 1.**
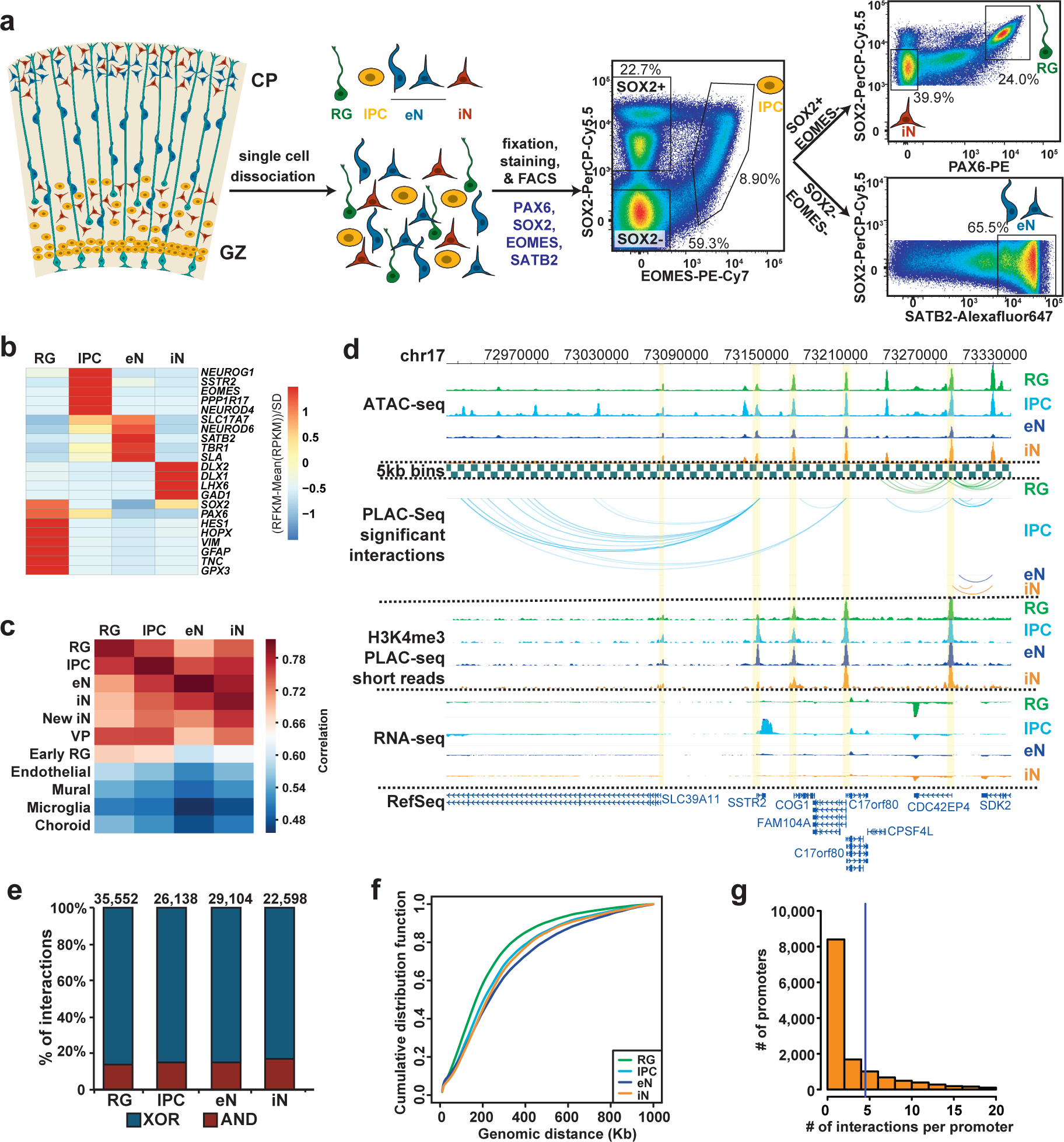
Experimental design and general features of the 3D epigenomic landscape during corticogenesis. (**a**) Schematic of the sorting strategy. Within the dorsal cortex, the germinal zone (GZ) is populated by radial glia (RG), which extend fibers towards the cortical plate (CP). These RG divide asymmetrically to produce intermediate progenitor cells (IPCs), which differentiate into excitatory neurons (eNs) that migrate along RG fibers towards the CP. At the same time, interneurons (iNs) can be found in both the GZ and CP. Microdissected GZ and CP samples were dissociated into single cells before being fixed, stained with antibodies for EOMES, SOX2, PAX6, and SATB2, and sorted using FACS. (**b**) Heatmap showing expression of key marker genes for RG, IPCs, eNs, and iNs. (**c**) Heatmap showing correlations between gene expression profiles for sorted cell populations and aggregate gene expression profiles from scRNA-seq datasets in the developing cortex. Cell types include newborn iNs from the medial ganglionic eminence (MGE), ventral progenitors including RG and IPCs from the MGE, microglia, and choroid plexus cells. (**d**) WashU Epigenome Browser snapshot of a 360 kb region (chr17:72,970,0000-73,330,000) showing IPC-specific chromatin interactions linked to *SSTR2* expression in IPCs. (e) Bar graph showing counts of MAPS interactions, with proportions of XOR (blue, only one interacting bin contains H3K4me3 peaks) and AND (red, both interacting bins contain H3K4me3 peaks) interactions displayed for each cell type. (**f**) Cumulative distribution function (CDF) plots showing interaction distances for each cell type. (**g**) Histogram showing the numbers of MAPS interactions at each promoter for each cell type.

The gene expression profiles of the sorted cell populations were both consistent with cellular identity and reproducible between individuals (**Fig. 1b and Supplementary Fig. 2a**). Sorted RG expressed *VIM*, *HES1*, *GPX3*, and *GFAP*, with little to no expression of marker genes for other cell types, whereas sorted IPCs expressed the IPC marker genes *EOMES*, *SSTR2*, and *NEUROD4*. In concordance with previous reports, *PAX6* was expressed in both RG and IPCs. Sorted eNs expressed the eN marker genes *SLA*, *SLC17A7*, *SATB2*, and *TBR1*, whereas *DLX1*, *DLX2*, and *GAD1* were exclusively detected in sorted iNs. When compared with aggregated scRNA-seq gene expression profiles^1^, our sorted cell populations exhibited the highest correlation with their corresponding subtypes while also showing reduced correlation with cells from the endothelial, mural, microglial, and choroid plexus lineages (**Fig. 1c**). Based on these results, we determined that our sorting strategy was robust and our sorted cell populations were suitable for additional epigenomic profiling.

### Characterization of 3D epigenomic landscapes during corticogenesis

We performed H3K4me3-centric proximity ligation-assisted ChIP-seq (PLAC-seq) to identify chromatin interactions at active promoters, assay for transposase-accessible chromatin using sequencing (ATAC-seq) to demarcate open chromatin regions, and RNA sequencing (RNA-seq) to profile transcriptomes in the sorted RG, IPCs, eNs, and iNs (**Fig. 1d and Supplementary Table 2**). We first confirmed the reproducibility of all PLAC-seq and ATAC-seq replicates (**Supplementary Fig. 2b and 2c**). Next, we applied the MAPS pipeline^13^ to call significant H3K4me3-mediated cis-regulatory chromatin interactions in merged replicates for each cell type at a resolution of 5 kb. We identified 35,552, 26,138, 29,104 and 22,598 MAPS interactions for RG, IPCs, eNs, and iNs, respectively, with approximately 85% of the interactions classified as anchor to non-anchor (XOR), and the remaining interactions classified as anchor to anchor (AND) (**Fig. 1e and Supplementary Fig. 3a and 3b**). The median interaction distance was between 170 kb to 230 kb for each cell type (**Fig. 1f**), and the majority of interactions occurred within TADs in GZ and CP tissues^10^ (**Supplementary Fig. 3c**).

**Figure 2.**
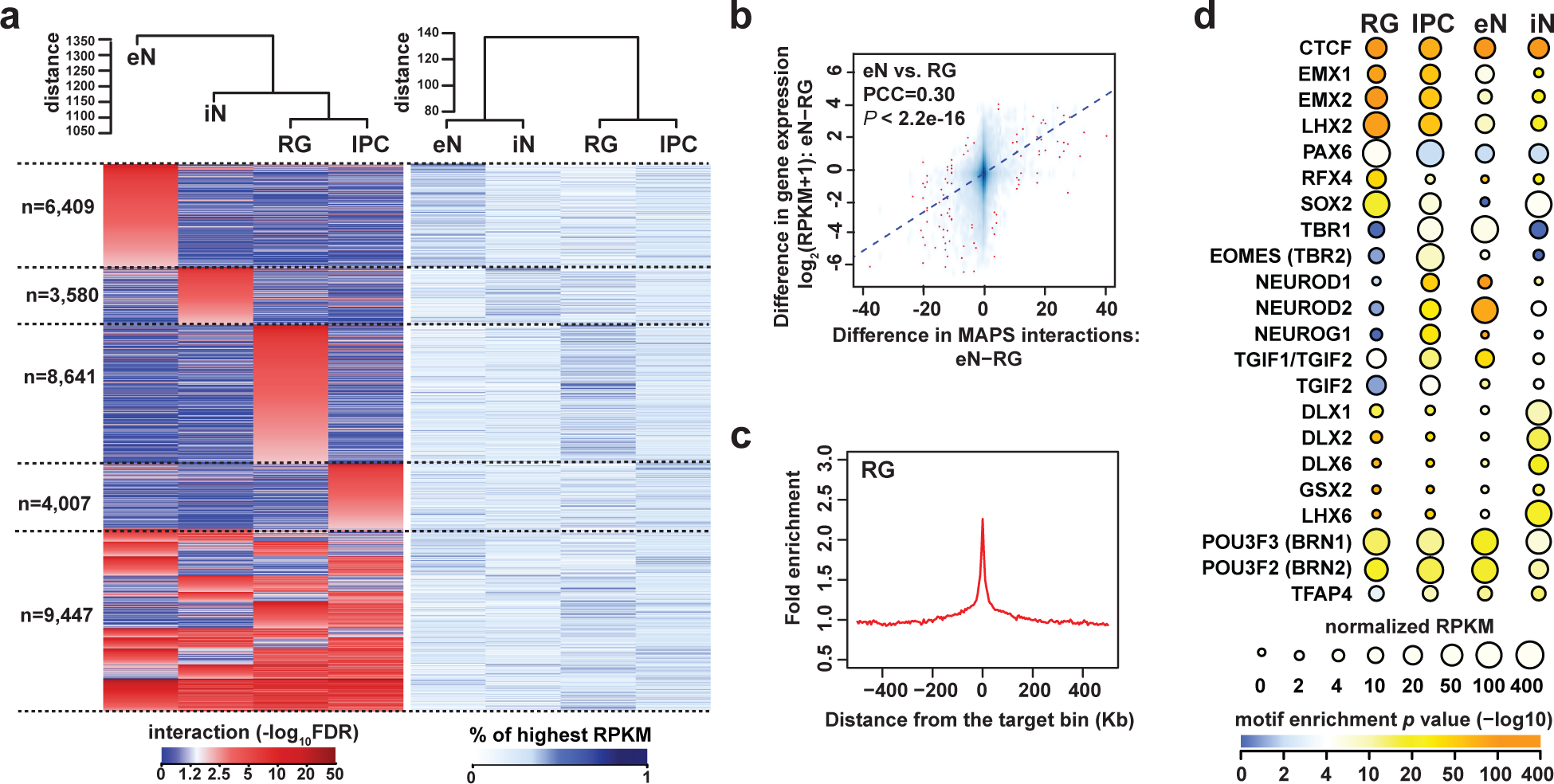
H3K4me3-mediated chromatin interactions contribute to cell type-specific gene regulation. (**a**) Heatmaps displaying interaction scores (left) and gene expression (right) for unique XOR interactions grouped according to their cell type specificity. Hierarchical clustering dendrograms for each heatmap are also shown (top). (**b**) Scatterplot showing positive correlation between the difference in the number of MAPS interactions at each promoter and the difference in expression of the corresponding genes between RG and eNs (Pearson product-moment correlation coefficient, two-sided t-test, *P* < 2.2×10^-16^). The fitted trendline based on linear regression is also shown. (**c**) Fold enrichment of open chromatin regions over distance-matched background regions in 1 Mb windows around distal interacting regions for MAPS interactions in RG. (**d**) Enrichment of TF motifs at open chromatin regions in cell type-specific interacting distal regions for each cell type. The color of each dot represents the degree of enrichment (-log10P-value) for each TF motif, and the size of each dot represents the gene expression of the corresponding TF.

**Figure 3.**
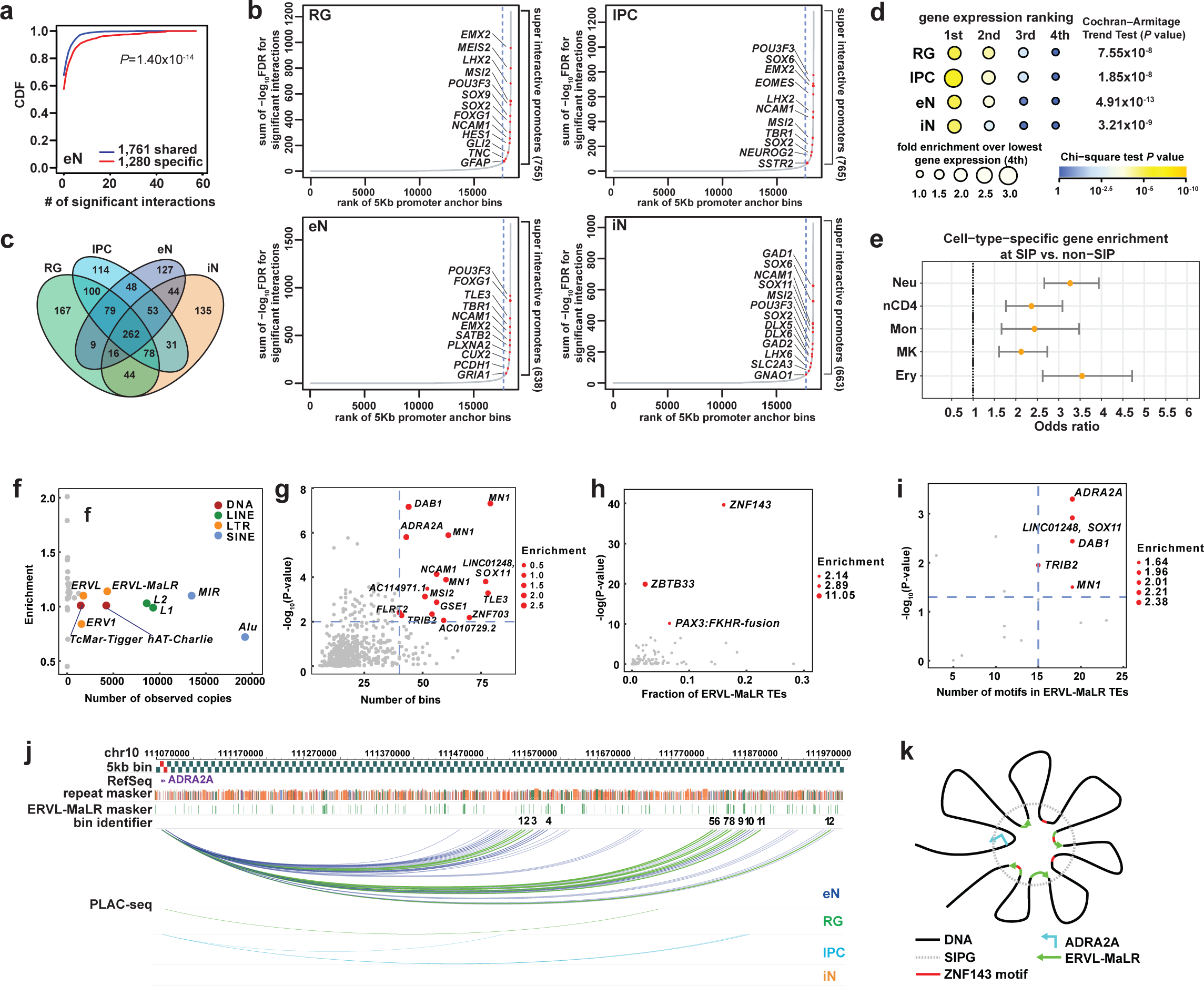
Super interactive promoters are enriched for lineage-specific genes. (**a**) CDF plots showing the numbers of MAPS interactions for shared versus cell type-specific genes in eNs (two sample t-test, two-sided, *P* = 1.40×10^-14^). (**b**) Plots showing the ranked cumulative interaction scores for 3D anchor bins in each cell type, defined as the sum of the -log10FDR for MAPS interactions coincident to each bin. Super interactive promoters (SIPs) are defined as promoters located to the right of the knee of each curve (dashed lines). Example SIPs, including those for lineage-specific genes, are highlighted for each cell type. (**c**) Venn diagram displaying the cell type-specificity of SIPs in RG, IPCs, eNs, and iNs. (**d**) The number of genes called as SIPs was divided by the total number of SIPs and non-SIPs for genes with the 1^st^, 2^nd^, 3^rd^, and 4^th^ highest expression among all four cell types. The fold enrichment was calculated relative to the group with the 4^th^ highest expression for each cell type. (**e**) Forrest plot showing that SIPs called in hematopoietic cells are also enriched for cell type-specific over shared genes. 95% confidence intervals are displayed. (**f**) Scatterplot showing both the enrichment and the number of observed copies of TE families in SIPGs for eNs. TE families occupying more than 1% of the genome are colored. (**g**) Scatterplot showing the enrichment of ERVL-MaLR TEs in SIPGs for eNs (hypergeometric P-value, see methods). SIPGs with 40 or more distal interacting bins and *P* < 0.01 are highlighted. (**h**) Scatterplot showing the enrichment of TF motifs at ERVL-MaLR TEs in SIPGs highlighted in (**g**). Only TF motifs with length > 12 bp are shown. (**i**) Scatterplot showing the enrichment of ZNF143 motifs at ERVL-MaLR TEs in SIPGs highlighted in (**g**) (Poisson distribution, see methods). ZNF143 motifs occurrences were detected using FIMO using a threshold of *P* = 0.0001. (**j**) WashU Epigenome Browser snapshot of the *ADRA2A* SIPG. MAPS interactions targeting the 12 distal interacting bins containing ERVL-MaLR-derived ZNF143 motifs are highlighted. (**k**) Potential mechanism for the contributions of TEs towards SIP formation.

### H3K4me3-mediated chromatin interactions contribute to cell type-specific gene expression

Since H3K4me3 is a histone mark associated with active promoters, we were interested to determine to what extent 3K4me3-mediated chromatin interactions influenced gene expression. First, we observed that the sorted cell populations clustered by developmental age based on their interaction strengths (**Fig. 2a**). This is consistent with iNs at this age possessing several characteristics of progenitor cells such as high *SOX2* expression (**Fig. 1a and 1b**). Genes participating in cell type-specific interactions are enriched for biological processes associated with their respective cell types, including cell proliferation for RG and IPCs, neuron projection development for IPCs and eNs, and synaptogenesis for eNs (**Supplementary Fig. 4a and Supplementary Table 3**). In addition, based on comparing interaction strength and gene expression side-by-side, we observe the two to be generally correlated (**Fig. 2a**). In fact, interaction strength and gene expression are globally correlated across all pairwise comparisons of cell types (**Fig. 2b and Supplementary Fig. 4b**), suggesting that gene expression is orchestrated by physical chromatin looping in a manner that is highly cell type-specific.

To investigate how chromatin interactions contribute to gene regulation in greater detail, we took advantage of the enrichment of open chromatin regions at distal interacting regions (**Fig. 2c and Supplementary Fig. 4c**) and performed transcription factor (TF) motif enrichment analysis using HOMER^14^ at the set of cell type-specific distal interacting regions for each cell type (**Fig. 2d and Supplementary Table 4**). PAX6, EOMES, and TBR1 are the most highly enriched motifs in RG, IPCs, and eNs, respectively, recapitulating their sequential expression along this developmental axis^15^. Meanwhile, motifs for progenitor-specific TFs including EMX1, EMX2, and LHX2 are enriched in RG and IPCs. The motif for RFX4, which was previously identified as an RG marker in the murine midbrain as well as the human telencephalon, is enriched in RGs^1,16^. Finally, the DLX1, DLX2, DLX6, GSX2, and LHX6 motifs are enriched in iNs, consistent with their roles in iN maturation and function^17-19^. Overall, our approach identifies both known and novel associations between TF binding at distal interacting regions and processes linked to cellular identity.

### Super interactive promoters are enriched for lineage-specific genes

The number of chromatin interactions at H3K4me3-marked promoters is only modestly correlated with gene expression (**Supplementary Fig. 5a**). One explanation is that individual genes are expressed to varying degrees in the context of their diverse cellular functions, and regulatory elements are better described as fine-tuning rather than independently inducing or silencing the expression of their cognate promoters. Multiple regulatory interactions can also exert synergistic or nonlinear effects on gene expression. To examine the relationship between gene expression and chromatin interactivity in greater detail, we first demonstrate that cell type-specific genes tend to have more interactions than shared genes across all cell types (**Fig. 3a and Supplementary Fig. 5b**). Next, by ranking promoter-containing anchor bins according to their cumulative interaction scores, we identify a subset of promoters with unusually high degrees of chromatin interactivity, which we term super interactive promoters (SIPs) (**Fig. 3b**). In total, we annotate 755, 765, 638, and 663 SIPs in RG, IPCs, eNs, and iNs, respectively (**Fig. 3c and Supplementary Table 5**). SIPs are enriched for key lineage-specific genes including *GFAP* and *HES1* for RG, *EOMES* for IPCs, *SATB2* for eNs, and *GAD1*, *GAD2*, *DLX5*, *DLX6*, and *LHX6* for iNs. SIPs are also frequently shared across multiple cell types. For example, we identify SIPs for *FOXG1* and *POU3F3* (*BRN1*) in all four cell types, *SOX2* in the progenitor-like RG, IPCs, and iNs, and *TBR1* in the eN-like IPCs and eNs. Interestingly, a large number of promoters for lincRNA genes including *LINC00461* and *LINC01551* are annotated as SIPs, consistent with their patterns of expression in the developing cortex^20^. Globally, SIPs are enriched for in cell types with the highest expression of their genes among all four cell types, supporting their putative roles in lineage specification (**Fig. 3d**). To assess whether SIPs are a general feature for other cell types, we expanded our analysis to hematopoietic lineages with published promoter capture Hi-C datasets^21^. Consistent with our results in brain cells, SIPs identified in neutrophils, naive CD4+ T cells, monocytes, megakaryocytes, and erythroblasts are also enriched for cell type-specific over shared genes (**Fig. 3e**). Based on these lines of evidence, SIPs may represent a general mechanism used by cells to maintain the precise and robust expression of key genes underlying cellular identity and function.

### Specific families of transposable elements are implicated in SIP formation

Given the important roles SIPs may harbor in establishing cellular identity, we were interested in exploring potential mechanisms underlying their formation and evolution. Towards this goal, we evaluated the contribution of transposable elements (TEs), which are capable of propagating regulatory elements across the genome and influencing 3D genome architecture^22-24^. First, we analyzed the enrichment of TEs at the class, family, and subfamily levels in sequences defined by the union of SIPs and their distal interacting regions (SIP groups or SIPGs) (**Fig. 3f and Supplementary Fig. 6a-c**). Notably, the ERVL-MaLR family and many of its subfamilies are enriched in SIPGs across all four cell types. Since we detected the strongest enrichment of this family of TEs in eNs, we decided to focus on this particular lineage. In total, we identified 16 SIPGs in eNs that are statistically enriched for ERVL-MaLR TEs (hypergeometric test, *P* < 0.01) (**Fig. 3g**). Next, we used HOMER to perform TF motif enrichment analysis at ERVL-MaLR TEs within these 16 SIPGs and determined ZNF143 to be the most enriched motif (**Fig. 3h**). ZNF143 is an architectural protein which has previously been reported to mediate looping between promoters and distal regulatory elements^25^. Moreover, certain subfamilies of ERVL-MaLR TEs have been demonstrated to contribute to ZNF143 binding in 3T3 and HeLa cells^26^. The *ADRA2A* SIPG in eNs exhibited the highest enrichment of ERVL-MaLR-derived ZNF143 motifs (hypergeometric test, *P=*1.59×10^-6^) (**Fig. 3i**) and was associated with elevated *ADRA2A* expression in eNs (**Supplementary Fig. 6d**). It spans 42 distal interacting regions, 25 of which contain ERVL-MaLR TEs, and 12 of which contain one or more ERVL-MaLR-derived ZNF143 motifs (**Fig. 3j and Supplementary Fig. 6e**). Furthermore, ZNF143 motifs in TEs from multiple ERVL-MaLR subfamilies (THE1A, THE1C, MSTA) within the SIPG can be mapped back to ZNF143 motifs in the consensus sequences of the same subfamilies (**Supplementary Fig. 6f**). This supports a model in which ZNF143 motifs are coordinately expanded by ERVL-MaLR TE insertion, leading to increased binding site redundancy and strengthened assembly of the *ADRA2A* SIPG regulatory unit (**Fig. 3k**). Our results imply that TEs are capable of mediating the formation of higher order epigenomic features including SIPs^27^.

### Investigating features of the developmental trajectory from RG to eNs during corticogenesis

Since RG, IPCs, and eNs represent a developmental trajectory from dorsal cortex progenitors to mature functional neurons, we grouped genes according to their expression and cumulative interaction scores along this axis and identified groups corresponding to RG, IPCs, and eNs (groups 1-3) that are enriched for lineage-defining genes and biological processes (**Fig. 4a, Supplementary Fig. 7, and Supplementary Table 6**). We also identified groups with decreasing expression and increasing chromatin interactivity (group 4) as well as increasing expression and decreasing chromatin interactivity (group 5) from RG to eNs, which could represent late-silenced and early-silenced genes, respectively. Late-silenced genes are enriched for chromatin remodeling and epigenetic regulation terms, whereas early-silenced genes are enriched for eN-specific signatures. These results demonstrate that gene expression can be mediated by distinct modes of chromatin-mediated regulation during development.

**Figure 4.**
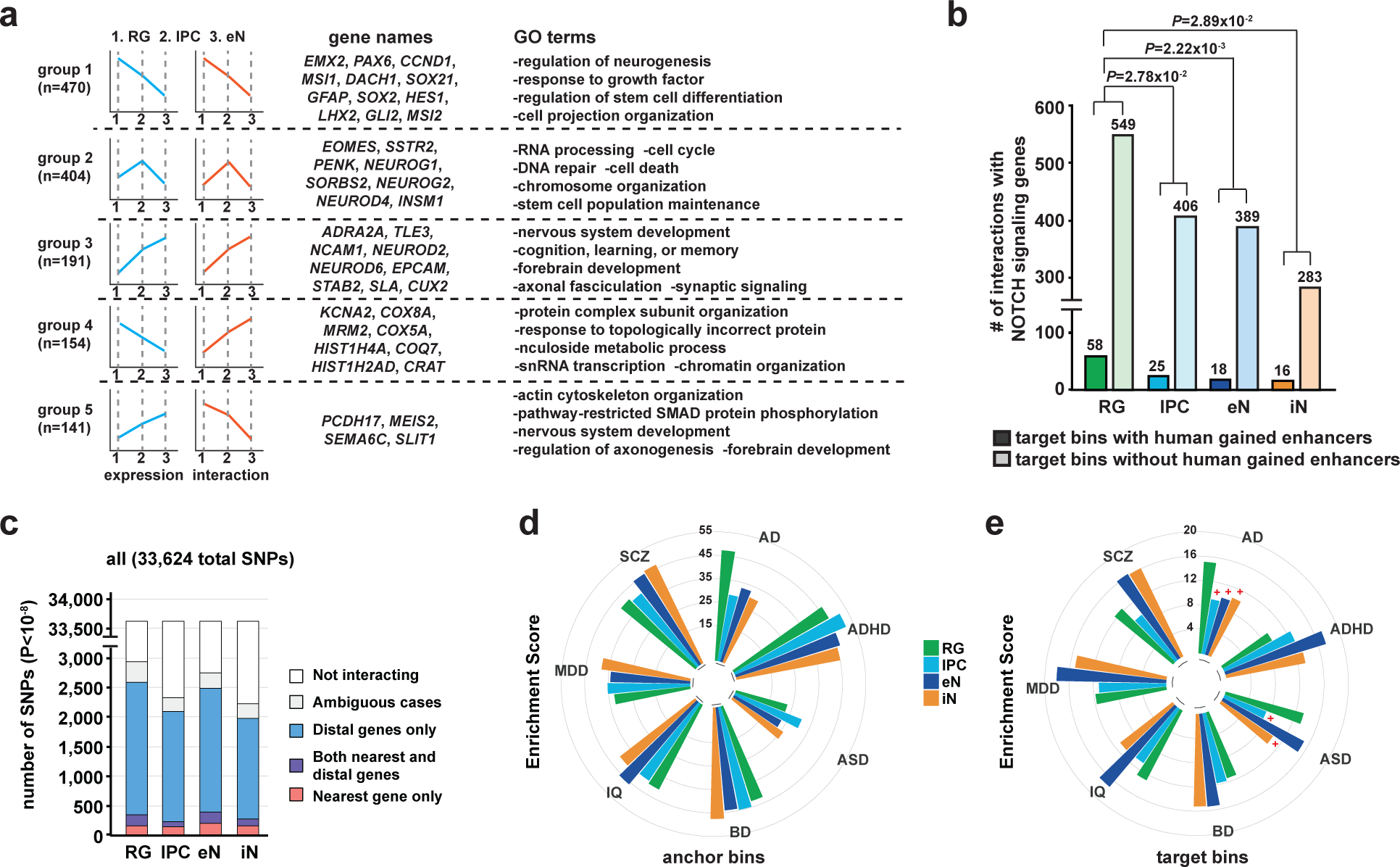
Investigating developmental trajectories during corticogenesis and partitioning heritability for complex neuropsychiatric disorders and traits. (**a**) Gene groups identified based on their changes in expression and chromatin interactivity along the transition from RG to eNs. Group 1 represents stem cell genes with decreasing expression and chromatin interactivity from RG to eNs. Group 2 represents IPC-specific genes with the highest expression and chromatin interactivity at the IPC stage. Group 3 represents genes with increasing expression and chromatin interactivity from RG to eNs. Groups 4 and 5 are characterized by anti-correlated expression and chromatin interactivity and may represent late-silenced and early-silenced genes, respectively. Representative genes and GO terms are shown for each group. (**b**) Bar graph showing the numbers of MAPS interactions at Notch signaling genes targeting bins with and without human gained enhancers in each cell type (Chi-square test). (**c**) Bar graph showing the numbers of unique GWAS SNPs (*P* < 10^-8^) interacting with their nearest gene only, with both their nearest and distal genes, or with distal genes only for each cell type across all neuropsychiatric traits. (**d-e**) LDSC enrichment scores for each neuropsychiatric trait and cell type, stratified by 3D anchor and target bins. Results with *P* > 0.05 are indicated.

Human corticogenesis is dramatically distinct from that in other mammalian species, driven in large part by the increased diversity and proliferative capacity of cortical progenitors during development which results in the increased size and complexity of the human brain^28^. In particular, Notch signaling genes have been implicated in the clonal expansion of RG, which constitute the major subtype of cortical progenitors in the cortex^29,30^. Here, we find that RG are enriched relative to other cell types for chromatin interactions at Notch signaling genes from the AmiGO database^31^ (**Fig. 4b**). Compared to other cell types, chromatin interactions in RG also target a significantly higher proportion of human gained enhancers identified through comparative analyses of H3K4me2 and H3K27ac ChIP-seq signal in human, rhesus macaque, and mouse brains^32^. Therefore, 3D epigenomic landscapes are capable of identifying lineage-specific pathways contributing to human-specific aspects of cortical development. In addition, we provide detailed annotations of gene targets for human gained enhancers and in vivo-validated enhancer elements from the Vista Enhancer Browser^33^ in **Supplementary Table 7**.

### Leveraging 3D epigenomic landscapes to partition heritability for complex neuropsychiatric disorder- and trait-associated variants

Chromatin interactions identified in our sorted cell populations represent a unique in vivo resource for mapping complex neuropsychiatric disorder- and trait-associated variants to their target genes (**Figure 4c and Supplementary Table 8**). They additionally enable the assessment of cell type-specific patterns of SNP heritability. To partition SNP heritability using our 3D epigenomic annotations, we employed linkage disequilibrium score regression (LDSC)^34,35^ using summary statistics from genome wide association studies (GWAS) for the following neuropsychiatric traits: Alzheimer’s disease (AD), attention deficit hyperactivity disorder (ADHD)^36^, autism spectrum disorder (ASD)^37^, bipolar disorder (BD)^38^, intelligence quotient (IQ)^39^, major depressive disorder (MDD)^40^, and schizophrenia (SCZ)^41^. Overall, we observed significant levels of heritability enrichment in 3D anchor bins for every cell type and neuropsychiatric trait we evaluated (15.13 < enrichment score < 51.57, 1.27×10^-40^ < *P* < 0.02) (**Fig. 4d**). These findings are largely expected as the majority of interacting promoters are shared across our cell types. When we restricted our analysis to 3D target bins, we observed dramatically distinct patterns of cell type-specific heritability enrichment (**Fig. 4e**). For example, ASD SNP heritability was significantly enriched for only in RG and eNs (*P* = 1.07×10^-3^ and 4.75×10^-4^, respectively), and AD SNP heritability was significantly enriched for only in RG (*P* = 3.99×10^-6^). Our findings reflect the cell type-specific nature of distal regulatory elements that are dysregulated during disease and underscore the importance of leveraging 3D epigenomic annotations to interpret variants that are located in non-coding regions of the genome.

### Functional characterization of enhancers in primary cells using CRISPRview

Validating distal regulatory elements in primary cells has historically been challenging, with most experiments to date performed using cell lines or iPSC-derived cells^10,42^. Here, we present CRISPRview, a novel approach combining CRISPRi, RNAscope, and immunostaining to validate enhancers in heterogeneous cultures of primary cells at the single cell level (**Fig. 5a**). We use CRISPRview to validate multiple enhancers in RG at the *GPX3*, *TNC*, and *HES1* loci, all of which harbor RG-specific chromatin interactions and are differentially expressed in RG (**Fig. 5b-d**). Furthermore, the promoters for *TNC* and *HES1* are annotated as SIPs in RG. First, sgRNAs were designed to target open chromatin regions physically interacting with the promoters of *GPX3*, *TNC*, and *HES1* (**Supplementary Table 9**). Next, primary cultures of microdissected and dissociated GZ samples between GW17 to GW19 were infected with lentivirus expressing the experimental sgRNA, dCas9-KRAB, and mCherry in combination with lentivirus expressing control sgRNA, dCas9-KRAB, and GFP. After five additional days in culture, the cells were fixed and stained using antibodies for mCherry, GFP, the RG marker GFAP, and RNAscope probes (ACD^TM^) targeting intronic regions of the genes of interest. Finally, high resolution images were taken using confocal microscopy, and the number of punctate dots representing individual nascent transcripts were compared between experimental (mCherry+) and control (GFP+) sgRNA-treated GFAP+ RG (**Fig. 5b-d**). The SMART-Q pipeline was specifically developed in our lab for image analysis (see attached manuscript, submitted).

**Figure 5.**
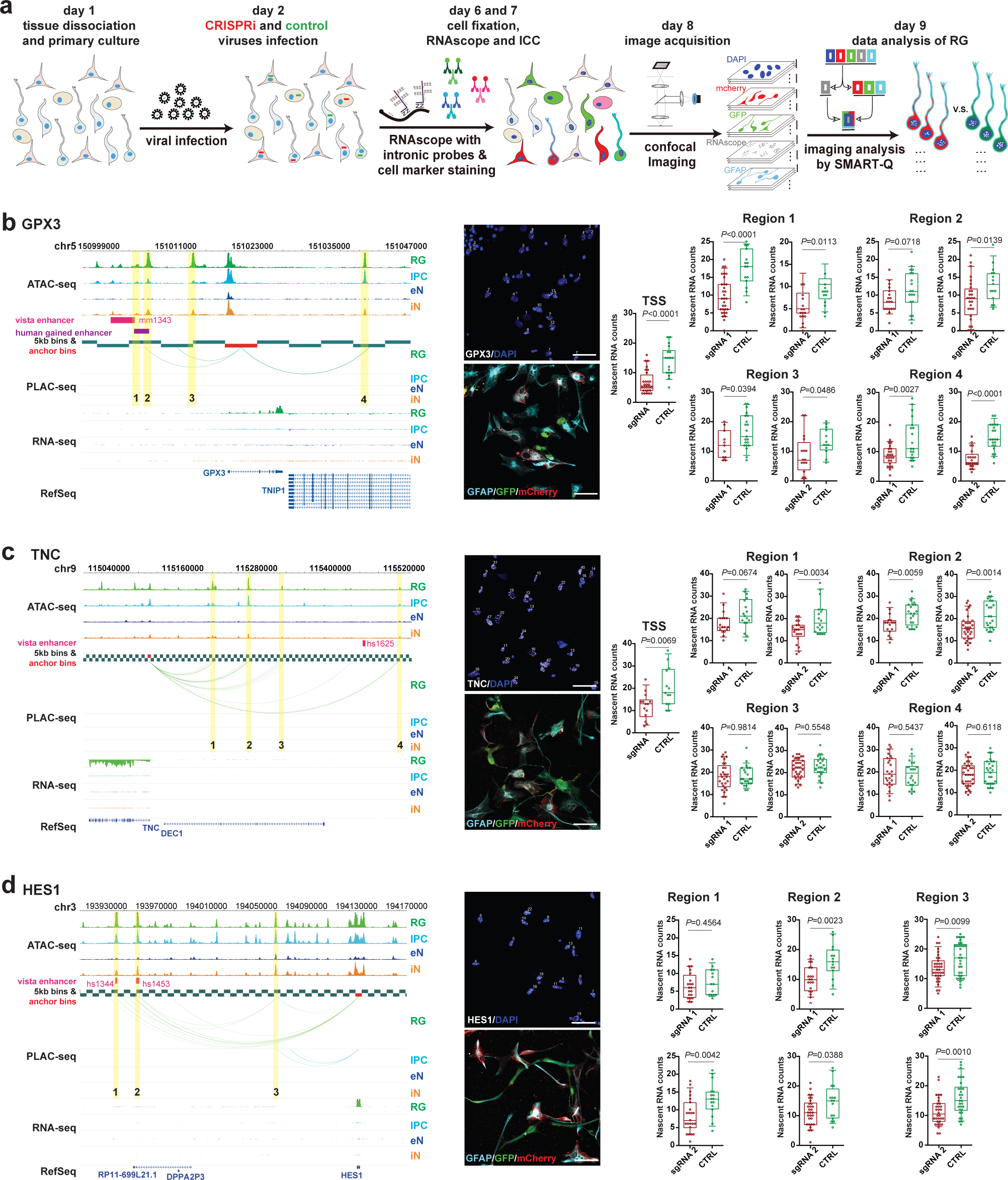
Functional characterization of distal interacting regions using CRISPRview. (**a**) CRISPRview workflow. Image analysis was performed using the SMART-Q pipeline. (**b**-**d**) Functional characterization of distal interacting regions at the *GPX3*, *TNC*, and *HES1* loci. For each locus, a WashU Epigenome Browser snapshot shows chromatin interaction bridging the promoters of *GPX3*, *TNC*, and *HES1* and distal interacting regions containing open chromatin regions (highlighted) which were targeted by sgRNAs for CRISPRi silencing. Representative images show staining for RNAscope probes targeting intronic regions for the genes of interest (white), DAPI (blue), the RG marker GFAP (light blue), mCherry (red), and GFP (green). The scale bar is 50 μm. Box plots show the results of CRISPRi silencing for each targeted region. The open circles represent single cells, and nascent transcript counts for experimental (mCherry+) versus control (GFP+) sgRNA-treated RG are represented on the y-axis (Student’s t-test, two-tailed). The median, upper and lower quartiles, and 10% to 90% range are indicated.

All four regions interacting with the *GPX3* promoter (regions 1-4) were found to exhibit downregulation of *GPX3* expression upon CRISPRi targeting (**Fig. 5b**). Notably, region 1 overlaps both a human gained enhancer^32^ and a Vista enhancer element (mm1343)^33^, supporting its function as an enhancer in RG. Next, we investigated the locus for *TNC*, a RG-specific gene implicated in neuronal migration, axon guidance, and synaptic plasticity.

We found that two of its interacting regions exhibited significant downregulation of *TNC* expression upon CRISPRi targeting (regions 1 and 2), but that silencing of its other two interacting regions did not result in notable changes in *TNC* expression (regions 3 and 4) (**Fig. 5c**). This could be due to the presence of alternative regulatory elements or structural interactions at these loci. Finally, HES1 is a lineage-defining TF for RG, and we positively validated all three regions that were found to interact with the *HES1* promoter (regions 1-3) (**Fig. 5d**). The observation of small but significant changes in gene expression upon CRISPRi targeting supports the hypothesis that multiple interactions work in concert to titrate the expression of key genes linked to cellular identity. In addition, the observed broad distributions of nascent transcript counts likely reflects the stochastic nature of transcription in single cells, stressing the importance of employing an approach combining resolution, sensitivity, and cell type-specificity for validating enhancers in single cells.

## Discussion

Recent publications leveraging single cell sequencing have highlighted the heterogeneity of the developing human cortex, underscoring the necessity of studying epigenomic regulation in a cell type-specific manner. Within the dorsal cortex alone, there are a massive variety of cell types from multiple sources both within and outside the developing neural tube, including RG, IPCs, eNs, MGE-derived iNs, CGE-derived iNs, microglia, endothelial cells, and subplate neurons. Despite large differences in maturation state and lineage, many of these cell types share intriguing similarities in terms of gene expression. For example, iNs express a number of TFs typically associated with RG proliferation such as *SOX2* or eN differentiation such as *ASCL1* and *NPAS3*^1^. Therefore, bulk measurements cannot reliably reveal the nuanced epigenomic programs driving gene expression in each cell type. By profiling 3D epigenomic landscapes in specific cell populations during corticogenesis, we not only demonstrate that gene regulation is closely linked to chromatin interactivity, we also identify SIPs that are highly cell type-specific and enriched for key lineage-specific genes. We uncover a potential mechanism by which specific families of TEs propagate binding sites for architectural proteins, facilitating the formation of multi-interaction clusters which may serve to sustain gene expression. While the analysis of TEs is currently constrained by the list of known motifs and the resolution of chromatin interactions identified in this study, future advances will help us further elucidate the contribution of TEs to 3D genome architecture and transcriptional regulation.

Cortical progenitors, eNs, and iNs are highly divergent in terms of their diversity, proliferative capacity, distribution, and functional characteristics between humans and mice^28^. Therefore, processes occurring during human cortex development cannot be fully recapitulated with mouse models. These non-murine features also indicate that enhancer mutations in humans may not adequately phenocopy to mice. Our dataset provides a comprehensive catalog of annotations for human gained enhancers and complex neuropsychiatric disorder- or trait-associated variants in cell types that are intricately tied to human cortex development, enabling the interpretation and prioritization of regulatory sequences for follow-up studies.

Finally, there is a need to perform cell type-specific validation for regulatory sequences in primary cells, especially as our understanding of epigenomic regulation matures over time. By combining immunostaining for cellular markers with the quantification of nascent transcripts in the nucleus, CRISPRview offers exquisite sensitivity and resolution for detecting cell type-specific changes in gene expression in single cells. Here, we used CRISPRview to successfully validate multiple regulatory elements in RG and observed subtle but significant changes in gene expression at the *GPX3*, *TNC*, and *HES1* loci. Further experiments leveraging CRISPRview in live tissue cultures should continue to reveal novel regulatory logic in a manner that is truly representative of the complex in vivo environment that is present during human cortex development.

## Acknowledgements

This work was supported by the UCSF Weill Institute for Neuroscience Innovation Award (to Y.S. and A.R.K.), the National Institutes of Health (NIH) grants R01AG057497, R01EY027789 and UM1HG009402 (to Y.S.), R35NS097305 (to A.R.K.), the Hillblom Foundation, and the American Federation for Aging Research New Investigator Award in Alzheimer’s Disease (to Y.S). This work was also supported by NIH grants R01HL129132, U544HD079124, and R01MH106611 (to Y.L.), R01HG007175, U24ES026699, and U01HG009391 (to T.W.), and the American Cancer Society grant RSG-14-049-01-DMC (to T.W). M.S. is supported by T32GM007175. M.P. is supported by the National Science Foundation Graduate Research Fellowship grant number 1650113. U.C.E. is supported by 5T32GM007618-42. This work was made possible in part by NIH grants P30EY002162 to the UCSF Core Grant for Vision Research, P30DK063720, and S101S10OD021822-01 to the UCSF Parnassus Flow Cytometry Core.

## Author contributions

Y.S., A.A.P., and A.R.K. conceived the study. Y.S., M.H., A.A.P., A.R.K. supervised the study. M.S., M.P., X.Y., I.R.J., X.C., U.C.E., and L.M. performed experiments. M.S., A.A., S.B., J.D.R., B.L., I.J., and M.H. performed computational analysis. C.F. and M.N.C. performed transposable element analysis under the supervision of T.W. J.W. and W.L. performed SNP heritability analysis under the supervision of Y.L. M.S., M.P., X.Y., and Y.S. analyzed and interpreted the data. Y.S., M.S., M.P., X.Y., and M.H prepared the manuscript with input from all other authors.

## Competing interests statement

The authors declare no competing financial interests.

## Code availability statement

A copy of the custom code used for data analysis and figure generation in this study is available upon request.

## Data availability statement

All datasets used in this study (PLAC-seq, ATAC-seq, RNA-seq) are available at the Neuroscience Multi-Omic Archive (NeMO Archive) under controlled access. Chromatin interactions, open chromatin regions, and gene expression results for each cell type can be accessed from the NeMO Archive using the following link: https://assets.nemoarchive.org/dat-uioqy8b Data can also be visualized on the WashU Epigenome Browser using the following link: http://epigenomegateway.wustl.edu/legacy/?genome=hg38&session=OCzw03b5lz&statusId=1958712809

## Methods

### Ethics statement

Deidentified embryonic brain tissue samples were collected with prior patient consent in strict observance of legal and institutional ethical regulations. All protocols were approved by the Human Gamete, Embryo, and Stem Cell Research Committee (GESCR) and Institutional Review Board at the University of California, San Francisco.

### Tissue dissociation

The tissue dissociation protocol was adapted from Nowakowski et al, 2017^1^. Briefly, samples were cut into small pieces in artificial cerebrospinal fluid before being added into pre-warmed papain dissociation media (Worthington #LK003150). The dissociation solution was incubated for 45 minutes at 37°C. Excess solution was removed and replaced with cell culture media. The pieces of tissue were triturated, filtered through a 70 µM nylon mesh, and centrifuged at 300 g for 8 minutes. The supernatant was removed and replaced with fresh culture media.

### Sample fixation

Mid-gestational human brain samples between GW15 and GW22 were fixed in 2% paraformaldehyde prepared in PBS with gentle agitation for 10 minutes at room temperature. Glycine was added to a final concentration of 200 mM to quench the reactions, and the samples were centrifuged at 500 g for 5 minutes at 4°C. The samples were washed twice with PBS before being frozen at -80°C until further processing.

### Permeabilization and staining

The cell pellet was thawed on ice and resuspended in PBS with 0.1% Triton-X-100 for 15 minutes. The cells were then washed twice with PBS and resuspended in 5% BSA in PBS for staining. Staining proceeded for at least one hour with FcR Blocking Reagent (Miltenyi Biotech, 1/20 dilution), EOMES-PE-Cy7 (Invitrogen, WD1928, 1/10 dilution), PAX6-PE (BD Biosciences, O18-1330, 1/10 dilution), SOX2-PerCP-Cy5.5 (BD Biosciences, O38-678, 1/10 dilution), and SATB2-Alexa Fluor 647 (Abcam, EPNCIR130A, 1/100 dilution). Following staining, the cells were centrifuged at 500 g for 5 minutes. The supernatant was removed, and the pellet was diluted into PBS. When sorting cells for RNA-seq libraries, 1% RNasin Plus RNase Inhibitor (Promega) was included in all buffers, PBS was prepared from RNase-free stocks, and acetylated RNase-free BSA was used to prepare 5% BSA in PBS for staining.

### FACS

AbC Total Antibody Compensation Beads (Thermo Fisher) were used to generate single color compensation controls prior to sorting. Sorting was conducted on either the FACSAria II, FACSAria IIu, or FACSAria Fusion instruments using a 70 µM nozzle, and cells were collected in 5 ml tubes pre-coated with FBS. A sample of each sorted population was reanalyzed on the same machine to assess purity. The cells were collected by centrifuging at 500 g for 10 minutes. The supernatant was removed, and the pellet was frozen at -80°C until further processing. When sorting cells for RNA-seq libraries, collection tubes were coated with both FBS and RNAlater (Thermo Fisher)

### Primary cell culture

Following dissociation, cells were plated onto Matrigel-coated coverslips in 48 well plates at a density of approximately 0.7×10^6^ cells per well. The cells were infected with lentivirus 24 hours after plating, and media was changed every two days. Media was composed of 96% DMEM/F-12 with GlutaMAX, 1% N-2, 1% B-27, and 1% penicillin/streptomycin. The cells were grown in 8% oxygen and 5% carbon dioxide, and they were harvested for fixation four days post-infection.

### PLAC-seq

PLAC-seq was performed according to the protocol from Fang et al., 2016^43^. 1 to 5 million cells were used to prepare each library. Digestion was performed using 100 U MboI for 2 hours at 37°C, and chromatin immunoprecipitation was performed using Dynabeads M-280 sheep anti-rabbit IgG (Invitrogen #11203D) mixed with 5 µg anti-H3K4me3 antibody (Millipore 04-745). TruSeq sequencing adapters were added during PCR amplification. Libraries were sent for paired-end sequencing on the HiSeq X Ten or NovoSeq 6000 instruments (150 bp paired-end reads). fastp was applied to trim reads to 100 bp, and replicates were merged and downsampled to normalize the number of usable reads before processing with MAPS.

### MAPS interaction calling

We used the MAPS pipeline to call significant long-range chromatin interactions from our PLAC-seq datasets. First, bwa mem was used to map raw reads to hg38. Unmapped reads and reads with low mapping quality were discarded, and the resulting filtered read pairs were processed as previously reported^13^. Briefly, we divided the genome into 5 kb bins and counted the number of read pairs representing interactions between 5 kb bins. To define our 3D anchor bins, we took the union of peaks called using MACS2 from read pairs with interaction distances < 1 kb for each cell type (1D H3K4me3 peaks). Based on this annotation, we classified interactions into AND, XOR, and NOT sets based on whether both, only one, or none of the interacting 5 kb bins overlapped 1D H3K4me3 peaks (**Supplementary Fig. 3a**). Since we were interested in identifying significant H3K4me3-mediated chromatin interactions, we retained only interactions in the AND and XOR sets for downstream processing. We also retained only intrachromosomal interactions with interaction distances between 10 kb and 1 Mb. These two criteria constituted our definition of usable reads.

For calling interactions, we applied a Poisson regression-based approach to normalize systematic biases from restriction sites, GC content, sequence repetitiveness, and ChIP enrichment. We fitted models for interactions in the AND and XOR sets separately and calculated false discovery rates (FDRs) for interactions based on their expected and observed contact frequencies between 5 kb bin pairs. Furthermore, we grouped interactions that were located within 15 kb of each other at both ends into clusters and classified all other interactions as singletons. To define our sets of significant long-range chromatin interactions, we retained only interactions with 12 or more reads, normalized contact frequencies (defined as the ratio between the observed and expected contact frequencies) ≥ 2, and FDR < 0.01 for clusters and FDR < 0.0001 for singletons. This was based on the reasoning that more biologically meaningful interactions are likely to appear in clusters, and singletons are more likely to present false positives.

### MAPS reproducibility analysis

PCA plots were generated based on the normalized contact frequencies for 5 kb bin pairs from our PLAC-seq datasets. Specifically, we first extracted AND and XOR 5 kb bin pairs based on cell type-specific 1D H3K4me3 peaks for each of the 11 replicates. We next applied zero-truncated Poisson regression, adjusting for the same systematic biases as the MAPS pipeline. Again, we derived normalized contact frequencies based on the ratio between the observed and expected contact frequencies between 5 kb bin pairs, with the expected contact frequencies being the fitted values from the zero-truncated Poisson regression. Normalized contact frequencies were then log-transformed and merged across the 11 replicates. This quantile normalized merged data was used to generate the PCA plots. We restricted our analysis to 5 kb bin pairs within 300 kb or 600 kb windows for **Supplementary Fig. 2c**.

### ATAC-seq

ATAC-seq was performed as previously described using the Nextera DNA Library Prep Kit (Illumina #FC-121-1030). First, fixed cells were washed once with ice cold PBS containing 1x protease inhibitor before being resuspended in ice cold nuclei extraction buffer (10 mM Tris-HCl pH 7.5, 10 mM NaCl, 3 mM MgCl2, 0.1% Igepal CA630, and 1x protease inhibitor) for 5 minutes. Next, 50,000 cells were counted out, exchanged into 50 μL 1x Buffer TD, then incubated with 2.5 μL TDE1 enzyme for 45 minutes at 37°C with shaking. Following transposition, 150 µL reverse crosslinking solution (50 µL 1 M Tris pH 8.0, 100 µL 10% SDS, 2 µL 0.5 M EDTA, 10 µL 5 M NaCl, 800 µL water, and 2.5 µL 20 mg/mL Proteinase K) was added to each reaction, and the reactions were incubated at 65°C overnight. On the next day, DNA was purified using Qiagen MinElute spin columns, amplified using Nextera primers, then size-selected for fragments between 300 and 1000 bp using AMPure XP beads. Libraries were sent for paired-end sequencing on the NovaSeq 6000 instrument (150 bp paired-end reads). Raw reads were mapped to hg38 and processed using the ENCODE pipeline (https://github.com/kundajelab/atac_dnase_pipelines) running the default settings. All sequencing reads were trimmed to 50 bp prior to mapping. The sets of optimal naive overlap peaks for each cell type were used for further downstream analysis.

### RNA-seq

We extracted total RNA from the sorted cell populations using the RNAstorm™ FFPE RNA extraction kit (Cell Data Sciences #CD501) starting from 5×10^5^ to 1.5×10^6^ cells. The quality of the extracted RNA was checked by calculating the percentage of RNA fragments with size > 200 bp (DV200) from the Agilent 2100 Bioanalyzer. RNA samples with DV200 >= 40% were used for library construction. First, they were depleted of ribosomal RNA using the KAPA RNA HyperPrep Kit with RiboErase (HMR #KK8560). Next, the RNA was used for first and second strand synthesis, dA-tailing, and sequencing adapter ligation. The cDNA was cleaned up and TruSeq sequencing adapters were added via PCR amplification. Libraries were sent for paired-end sequencing on the NovaSeq 6000 instrument (150 bp paired-end reads). Raw reads were aligned to hg38 using STAR running the standard ENCODE parameters, and transcript quantification was performed in a strand-specific manner using RSEM with the GENCODE 29 annotation. The edgeR package in R was used to calculate TMM-normalized RPKM values for each gene based on the expected counts and gene lengths for each replicate as reported by RSEM. The mean gene expression across all replicates for each cell type was used for further downstream analysis.

### TF motif enrichment analysis

We took the set of open chromatin regions participating in cell type-specific XOR interactions for each cell type and used the sequences in 200 bp windows around the peak summits to perform motif enrichment analysis using HOMER running the default settings. The complete set of vertebrate motifs from the JASPAR database were used for detection. The “-float” option was specified to optimize the detection threshold, and the entire genome was used as a background. Entries with similar or identical consensus TF motif sequences were grouped for brevity.

### GO enrichment analysis

Protein coding and noncoding RNA genes from GENCODE 29 participating in cell type-specific XOR interactions were used for GO enrichment analysis. Only genes participating in interactions with promoter open chromatin regions on one end and distal open chromatin regions on the other end were used. A minimum normalized RPKM of 0.5 was used to filter out genes that were not significantly expressed the corresponding cell types, and the resulting gene lists were input into DAVID 6.8 running functional annotation clustering with the default settings and the “GOTERM_BP_ALL” ontology.

### SIP identification

We devised an approach similar to calling super-enhancers^44^ to identify super interactive promoters (SIPs) using our MAPS interactions for each cell type. Specifically, we started from 18,373 anchor bins containing 1D H3K4me3 peaks annotated in at least one of the sorted cell populations. For each anchor bin, we calculated the cumulative interaction score for all its coincident MAPS interactions. For anchor bins without any MAPS interactions, the cumulative interaction score was calculated to be zero. We then prepared plots of the ranked cumulative interaction scores for anchor bins in each cell type and defined SIPs as promoters located to the right of the knee of each curve.

### Defining cell type-specific versus shared genes

We classified each gene as cell type-specific or shared according to its Shannon entropy across all four cell types. Specifically, we first calculated the relative expression of each gene in each cell type, defined as a gene’s normalized RPKM in the cell type divided by the sum of the gene’s normalized RPKMs across all four cell types. Next, we calculated the Shannon entropy based the gene’s relative expression in each of the cell types. A cell type-specific gene is characterized by low entropy, while a shared gene is characterized by high entropy. We classified a gene as cell type-specific if met the following conditions: its entropy was < 0.01, its normalized RPKM was > 1 in that cell type, and its normalized RPKM was highest in that cell type among all four cell types. All other genes with normalized RPKM > 1 across every cell type were classified as shared.

### TE family and subfamily enrichment in SIPGs

A SIP and its distal interacting regions are considered to be a SIP group or SIPG. TE enrichment in SIPGs was evaluated as follows. The foreground enrichment was defined as the number of copies of TEs from a given family or subfamily overlapping SIPGs in each cell type. The background enrichment was defined as the number of copies of TEs overlapping all interacting 5 kb bins. The overall enrichment was defined as the foreground enrichment divided by the background enrichment multiplied by the fraction of interacting 5 kb bins belonging to SIPGs. At least 50% of a TE had to intersect a 5 kb bin for it to be considered to overlap the 5 kb bin.

### ERVL-MaLR TEs enrichment in specific SIPGs

For each SIPG, the foreground enrichment was defined as the number of distal interacting regions with one or more copies of an ERVL-MaLR TE. The background enrichment was calculated by randomly permuting the locations of the distal interacting regions and counting the number of permuted regions with one or more copies of an ERVL-MaLR TE. This was performed over 100 such permutations. The overall enrichment was defined as the foreground enrichment divided by the background enrichment. The one-tailed P-value for each SIPG was calculated using the hypergeometric distribution as follows: *P* = choose (m, q) x choose (n, k - q) / choose (m + n, k), where “q” is the number of 5 kb bins within the SIPG with one or more copies of an ERVL-MaLR TE, “m” is the number of 5 kb bins with one or more copies of an ERVL-MaLR TE on the same chromosome, “n” is the number of 5 kb bins with no copies of an ERVL-MaLR TE on the same chromosome, and “k” is the size of the SIPG.

### ZNF143 motif enrichment at ERVL-MaLR TEs in specific SIPGs

For each SIPG, the foreground enrichment was defined as the number of ZNF143 motifs occurring in ERVL-MaLR TEs as determined using FIMO in distal interacting regions for the SIPG. The background enrichment was defined as the number of ZNF143 motifs occurring in the SIPG, but not necessarily in the ERVL-MaLR-TEs. The overall enrichment was defined as the foreground enrichment divided by the background enrichment multiplied by fraction of the SIPG sequence occupied by ERVL-MaLR TEs. The one-tailed P-value was calculated using the Poisson distribution as follows: the number of events is the foreground enrichment, and the probability is the background enrichment multiplied by fraction of the SIPG sequence occupied by ERVL-MaLR TEs.

### Target gene annotation for enhancers and complex neuropsychiatric disorder- and trait-associated variants

To determine whether a human gained enhancer, Vista enhancer element, or GWAS SNP potentially interacted with a target gene, we determined whether any of its promoters participated in MAPS interactions with the feature of interest on the other end. All human gained enhancers and Vista enhancer elements were expanded to a width of 5 kb and all GWAS SNPs were expanded to a width of 1 kb to account for potential functional sequences around each feature. Furthermore, we determined the proportion of GWAS SNPs interacting with their nearest and/or distal genes, except when the promoters for the nearest gene and GWAS SNP fell within the same 5 kb bin and could not be resolved for MAPS interactions (“same fragment ambiguity”). We provide target gene annotations for human gained enhancers and Vista enhancer elements in **Supplementary Table 7** and GWAS SNPs in **Supplementary Table 8**. The overlap of each feature with open chromatin regions in each cell type is also reported.

### Partitioning heritability for complex neuropsychiatric disorder- and trait-associated variants

We employed stratified LD score regression^34,35^ to partition SNP heritability for neuropsychiatric traits using our cell type-specific 3D epigenomic annotations. Specifically, we first collected GWAS summary statistics for seven complex neuropsychiatric disorders and traits including Alzheimer’s disease (AD), attention deficit hyperactivity disorder (ADHD), autism spectrum disorder (ASD), bipolar disorder (BD), intelligence quotient (IQ), major depressive disorder (MDD), and schizophrenia (SCZ). We estimated the enrichment of SNP heritability for each complex neuropsychiatric disorder and trait separately based on 3D anchor or target bins from MAPS interactions for each cell type. 3D anchor bins contain H3K4me3 ChIP-seq peaks and are presumably enriched for active promoters, whereas 3D target bins are presumably enriched for distal regulatory elements such as enhancers.

### Validation of distal interacting regions using CRISPRview

The CRISPRi vector was modified from the Mosaic-seq^45^ and CROP-seq vectors^46^. The hU6-sgRNA expression cassette from the CROPseq-Guide-Puro vector (Addgene #86708) was cloned and inserted downstream of the WPRE element in the Lenti-dCas9-KRAB-blast vector (Addgene #89567). The blasticidin resistance gene was replaced with either mCherry or EGFP. sgRNAs targeting open chromatin regions in distal interaction regions were designed using CHOPCHOP. Single stranded DNA was annealed and ligated into the CRISPRi vector at the BsmBI cutting locus. Single clones were picked following transformation, and the sgRNA sequences were confirmed by Sanger sequencing. For lentiviral packaging, the CRISPRi vector, pMD2.G (Addgene #12259), and psPAX (Addgene #12260) were transformed into 293T cells using PolyJet (SignaGen Laboratories #SL100688) according to the manufacturer’s instructions. Virus-containing media was collected three times every 16 to 20 hours and concentrated using Amicon 10K columns. Collected lentivirus was stored immediately at -80°C. Primary cell cultures were infected with virus (MOI < 1) 24 hours after plating, and four days after infection, cells were harvested and fixed with 4% PFA for FISH and immunostaining.

FISH experiments detecting nascent transcripts were performed using the RNAScope Multiplex Fluorescent V2 Assay kit (ACDBio #323100) followed by immunostaining for cell type-specific markers. Probes targeting intronic regions for *GPX3* (ACDBio #572341), *TNC* (ACDBio #572361), and *HES1* (ACDBio #560881) were custom-designed, synthesized, and labeled with TSA Cyanine 5 (Perkin Elmer #NEL705A001KT, 1:1000 dilution). Next, fixed cells were pretreated with hydrogen peroxide for 10 minutes and Protease III for 15 minutes, and probes were hybridized and amplified according to the manufacturer’s instructions. Slides were washed with PBS before blocking with 5% donkey serum in PBS for 30 minutes at room temperature. Next, slides were incubated with primary antibodies against mCherry (Abcam ab205402), GFP (Abcam ab1218) and GFAP (Abcam ab7260) overnight at 4°C, followed by incubation with Alexa Fluor 488 donkey anti-mouse IgG (Thermo Fisher Scientific #A21202), Alexa-546 nm donkey anti-rabbit IgG (Thermo Fisher Scientific #A10040), or Alexa-594 nm goat anti-chicken IgG (Thermo Fisher Scientific #A11042) for 1 hour at room temperature. Three-dimensional confocal microscopy images were captured using a Leica TCS SP8 with a 40x oil-immersion objective lens (NA = 1.30). The z-step size was 0.4 µm. For five color multiplexed imaging, three sequential scans were performed to avoid overlapping spectra. The first excitation lasers were 405 nm and 594 nm, the second excitation lasers were 488 nm and 633 nm, and the third excitation laser was 561 nm. All images were obtained using the same acquisition settings. For FISH analysis, we developed an integrated Python-based pipeline called Single-Molecule Automatic RNA Transcription Quantification (SMART-Q) for quantifying nascent transcripts in single cells. Briefly, RNAscope signal was filtered then fitted in three dimensions using Gaussian models. Next, segmentation was performed on the DAPI channel in two dimensions to ascertain the location of each nucleus. Finally, segmentation was performed on the cell marker channel to identify RG-specific nuclei, and the positional RNAscope data was integrated with the segmentation results to determine the final quantification of nascent transcripts in each cell.

---

**Supplemental Figure 1.**
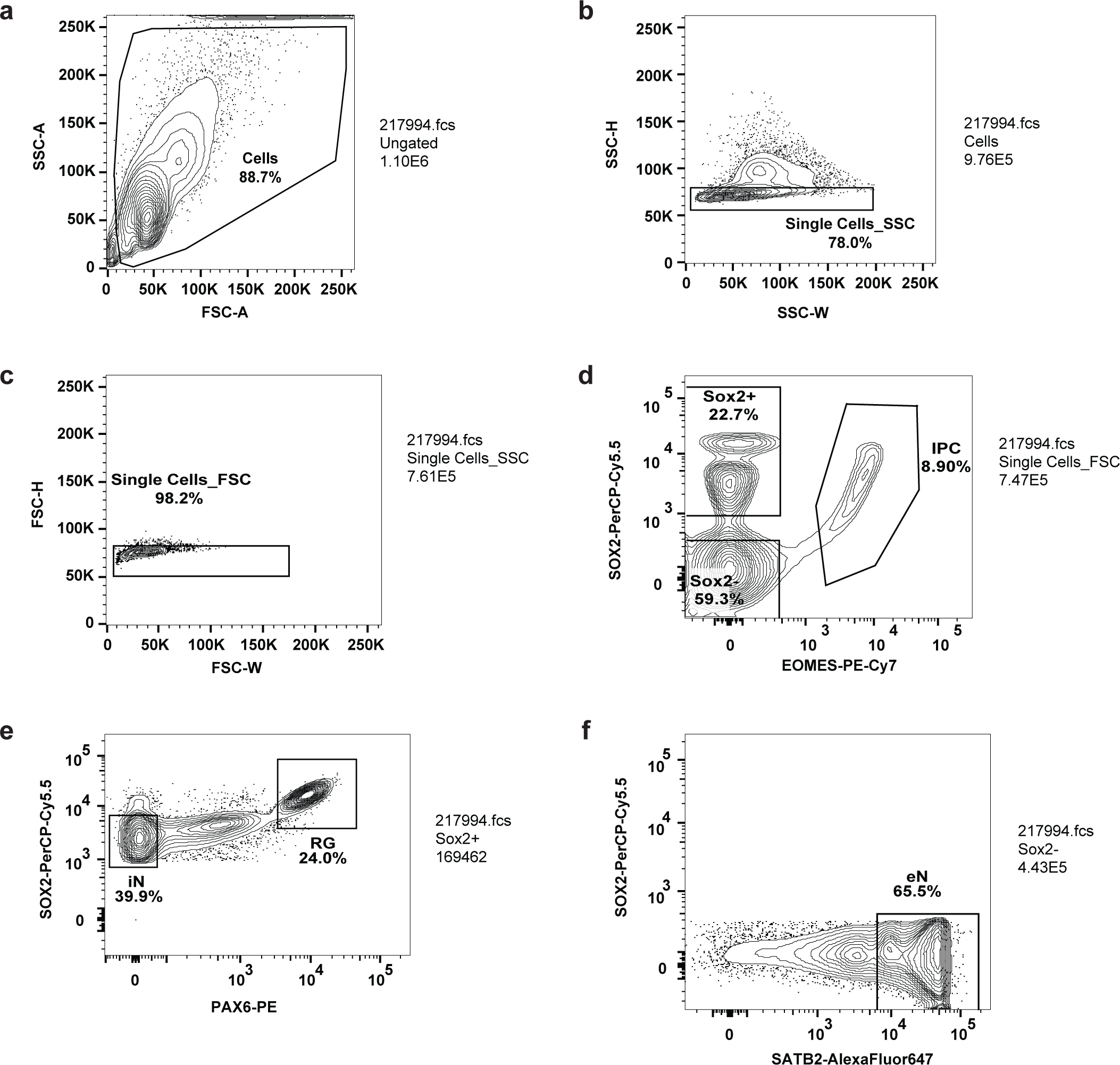
Representative contour plots depicting FACS gating strategy. (**a**) Cells were separated from debris of various sizes based on the forward scatter area (FSC-A) and side scatter area (SSC-A). Cells were then passed through two singlet gates using the width and height metrics of the (**b**) side scatter (SSC-H versus SSC-W) and (**c**) forward scatter (FSC-H versus FSC-W). (**d**) SOX2+, and SOX2-, and intermediate progenitor (IPC) populations were isolated by gating on EOMES-PE-Cy7 and SOX2-PerCP-Cy5.5 staining. (**e**) Radial glia (RG) and interneurons (iNs) were isolated as high PAX6/high SOX2 and medium SOX2/low PAX6 populations, respectively. (**f**) Excitatory neurons (eNs) were isolated from the SOX2-population by gating on SATB2-Alexa Fluor 647 staining.

**Supplementary Figure 2.**
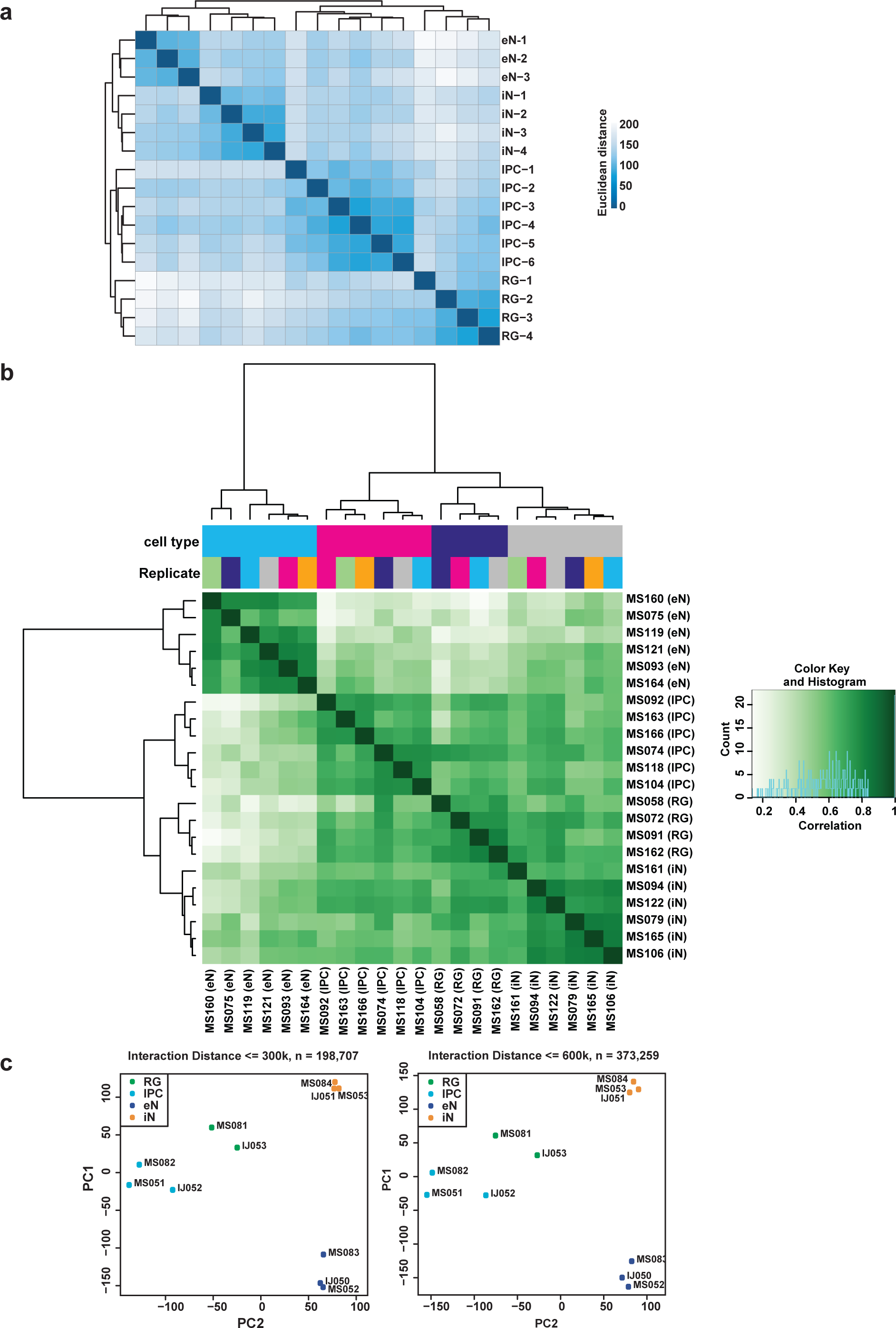
Reproducibility between replicates for RNA-seq, ATAC-seq, and PLAC-seq. (**a**) RNA-seq replicates were hierarchically clustered according to gene expression sample distances using DESeq2. (**b**) Heatmap with pairwise correlations and hierarchical clustering for read densities at the set of unified open chromatin regions for ATAC-seq replicates. (**c**) Principle component analysis (PCA) was performed based on the normalized contact frequencies across all PLAC-seq replicates (see methods). To assess the robustness of the results, we conducted the analysis separately for bin pairs within 300 and 600 kb interacting windows.

**Supplementary Figure 3.**
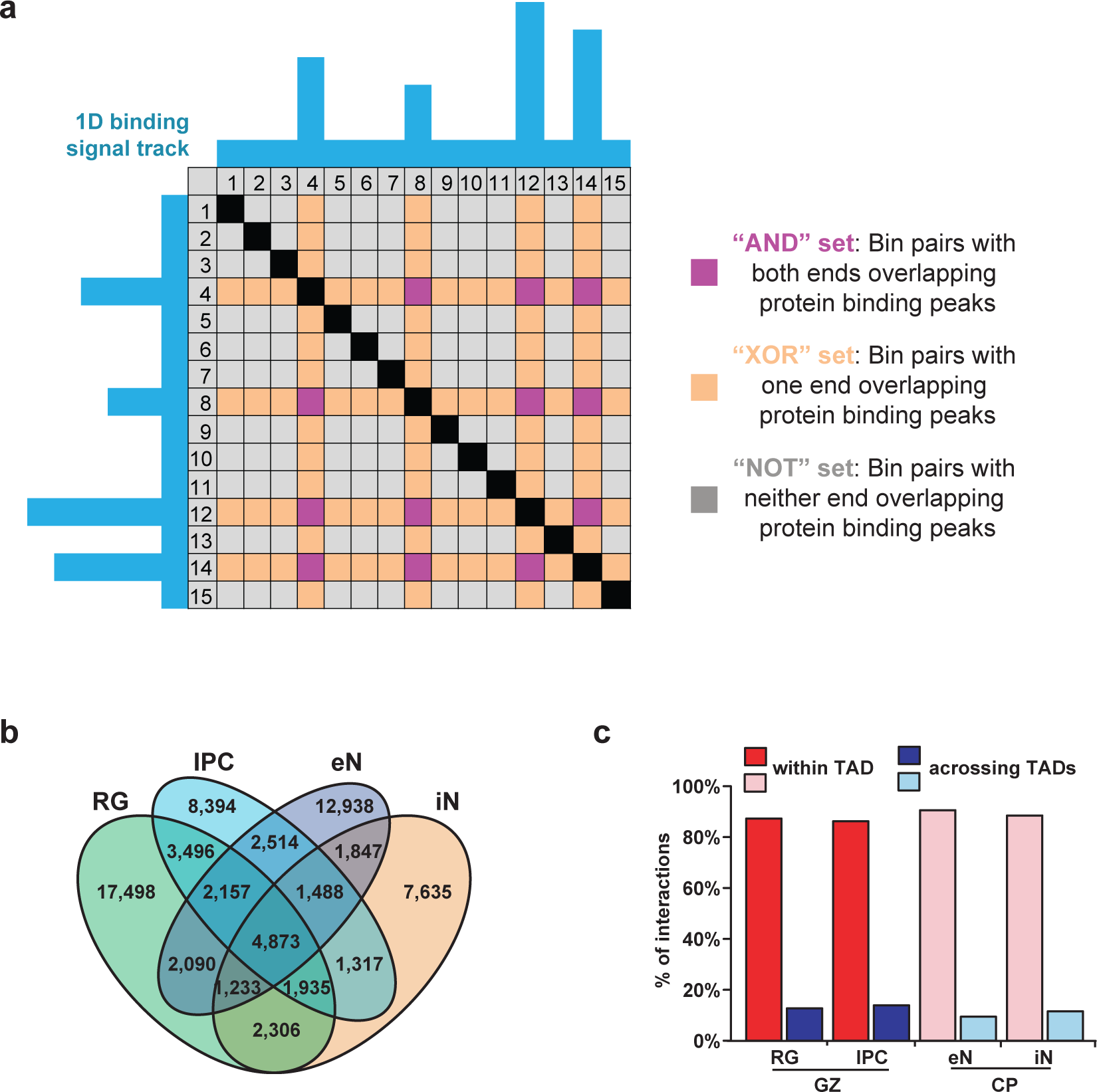
Identification of chromatin interactions using MAPS. (**a**) Illustration of AND and XOR sets in a representative PLAC-seq contact matrix. The blue tracks represent 1D H3K4me3 peaks at bin positions 4, 8, 12, and 14. The black cells represent interactions within the same bin. The purple cells represent interactions in the AND set where both of the interacting bins contain 1D H3K4me3 peaks. The orange cells represent interactions in the XOR set where only one of the interacting bins contains 1D H3K4me3 peaks. The grey cells represent interactions where neither of the interacting bins contains 1D H3K4me3 peaks. (**b**) Venn diagram displaying cell type-specificity of MAPS interactions for each cell type. (**c**) Proportions of MAPS interactions occurring within and across TADs in GZ and CP tissues for each cell type.

**Supplementary Figure 4.**
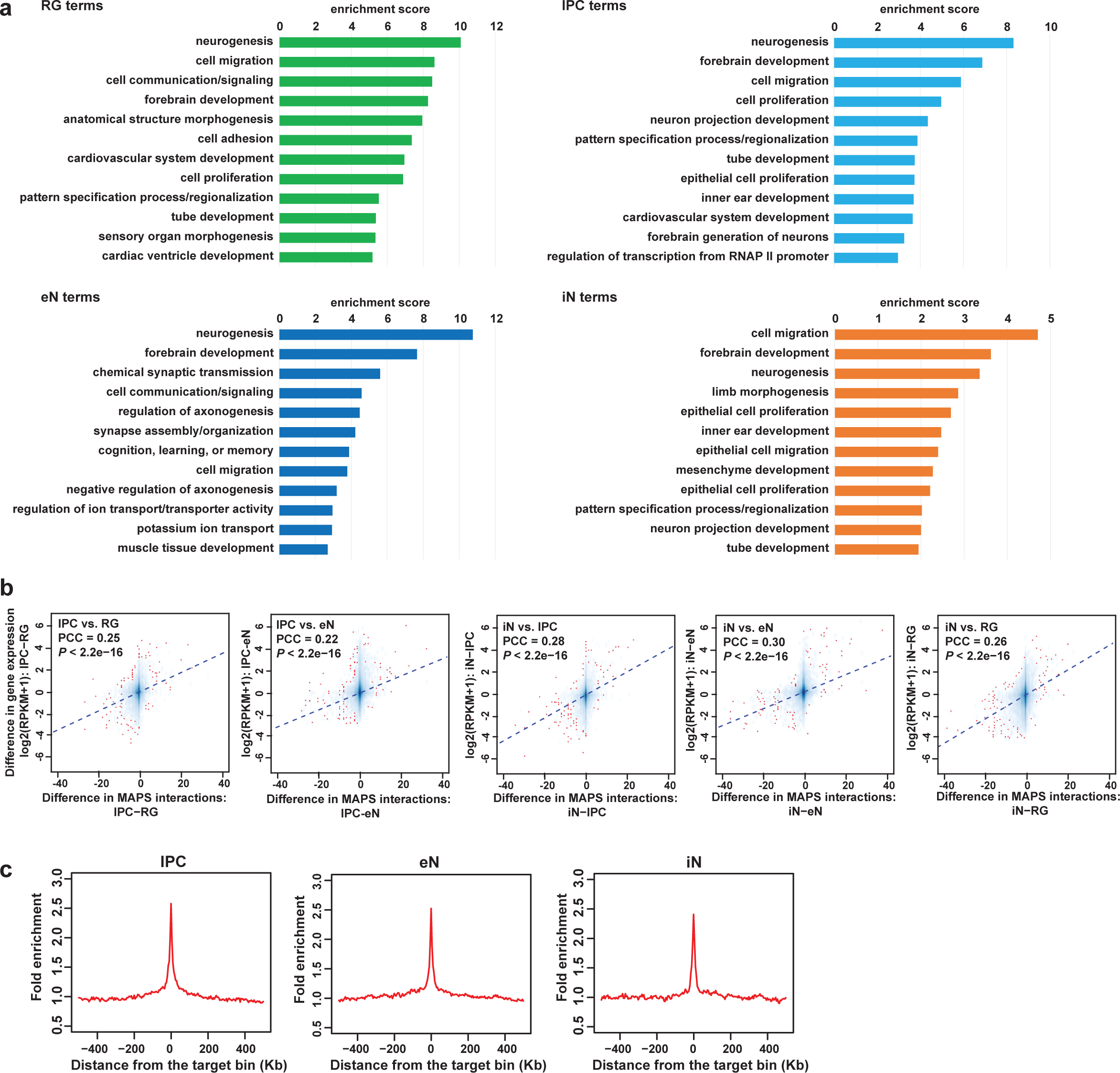
Contribution of 3D epigenomic landscapes to gene regulation. (**a**) GO enrichment analysis for genes whose promoters participate in cell type-specific interactions. The top annotation clusters from DAVID are reported along with their enrichment scores for each cell type. (**b**) Scatterplots showing positive correlation between the difference in the number of MAPS interactions at each promoter and the difference in expression of the corresponding genes between all pairs of cell types (Pearson product-moment correlation coefficient, two-sided t-test, *P* < 2.2×10^-16^ for all cell types). Fitted trendlines based on linear regression are also shown. (**c**) Fold enrichment of open chromatin regions over distance-matched background regions in 1 Mb windows around distal interacting regions for MAPS interactions in IPCs, eNs, and iNs.

**Supplementary Figure 5.**
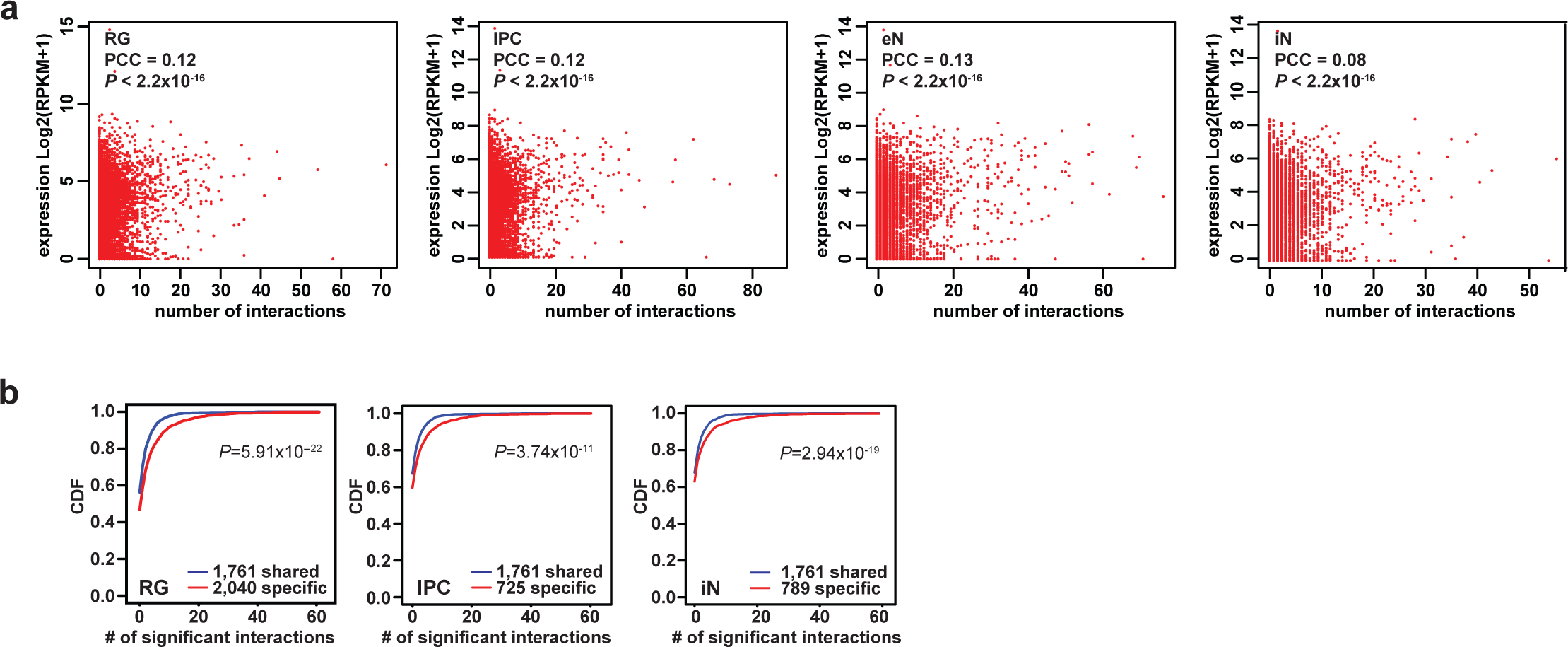
Correlations between chromatin interactions and gene expression for cell-type specific and shared genes. (**a**) Scatterplots showing the correlation between numbers of MAPS interactions and gene expression at promoters in each cell type. (**b**) Cumulative distribution function (CDF) plots showing the numbers of MAPS interactions for shared versus cell type-specific genes in RG, IPCs, and iNs (two sample t-test, two-sided, *P* = 5.91×10^-22^, 3.74×10^-11^, and 2.94×10^-19^ for RG, IPCs, and iNs, respectively).

**Supplementary Figure 6.**
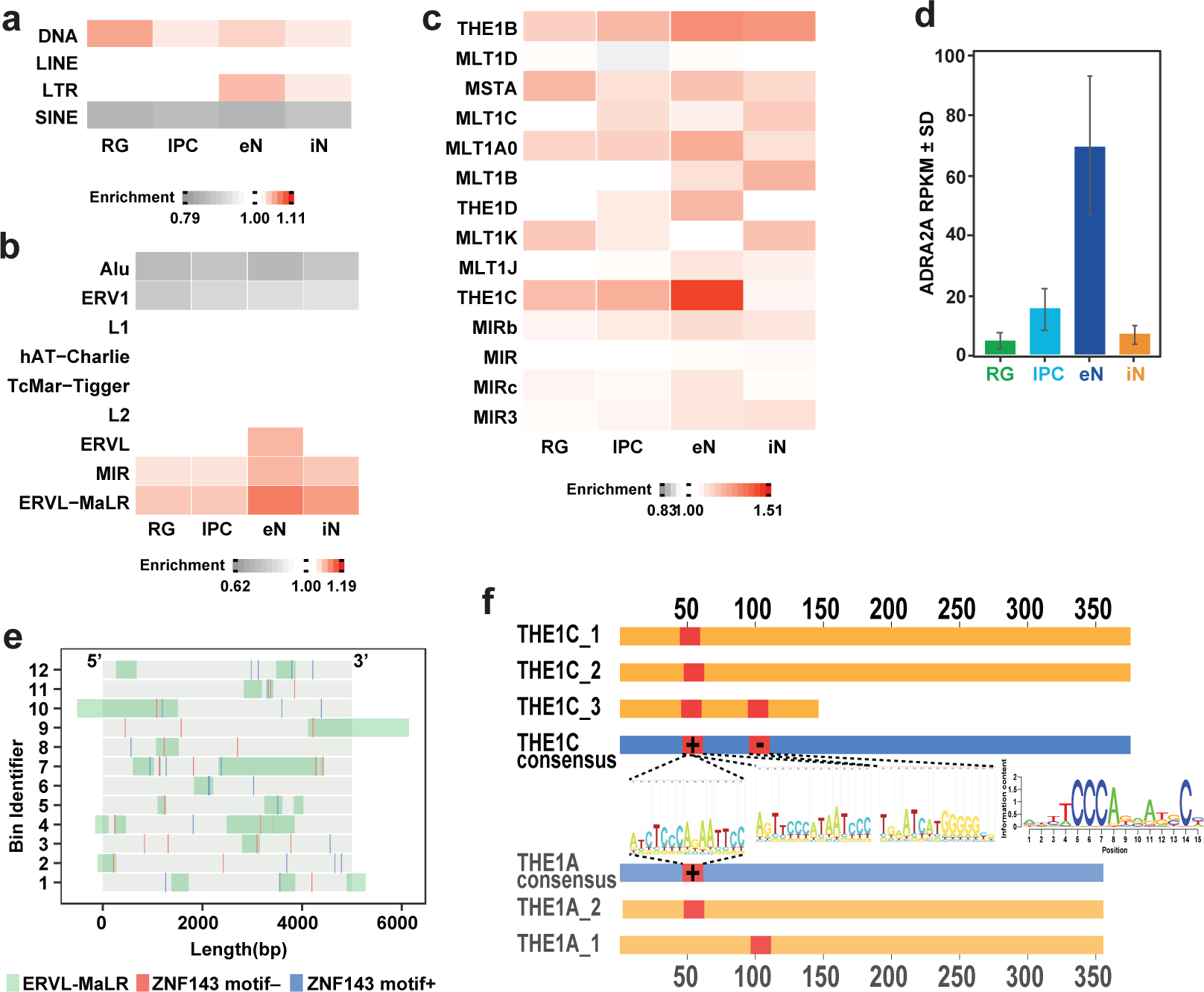
Specific families of transposable elements are implicated in SIP formation. **(a-c)** Enrichment of TEs in SIPGs at the class (**a**), family (**b**), and subfamily (**c**) levels for each cell type. Only families occupying more than 1% of the genome are shown in (**b**). Only subfamilies from the MIR and ERVL-MaLR families occupying more than 0.1% of the genome are shown in (**c**). (**d**) Bar graph shows elevated *ADRA2A* gene expression in eNs. (**e**) Illustration of the 12 distal interacting regions in the *ADRA2A* SIPG containing at least one ERVL-MaLR-derived ZNF143 motif. ZNF143 motifs are indicated and colored by strand. The bin identifier corresponds to the labels in **Fig. 3j**. (**f**) Illustration of the conservation of ZNF143 binding motifs in ERVL-MaLR TEs. Blue bars indicate consensus sequences, yellow bars indicate individual copies of ERVL-MaLR TEs in the *ADRA2A* SIPG, and red bars indicate ZNF143 motifs. The positions of the ZNF143 motifs relative to the ERVL-MaLR TE sequences was determined using FIMO.

**Supplementary Figure 7.**
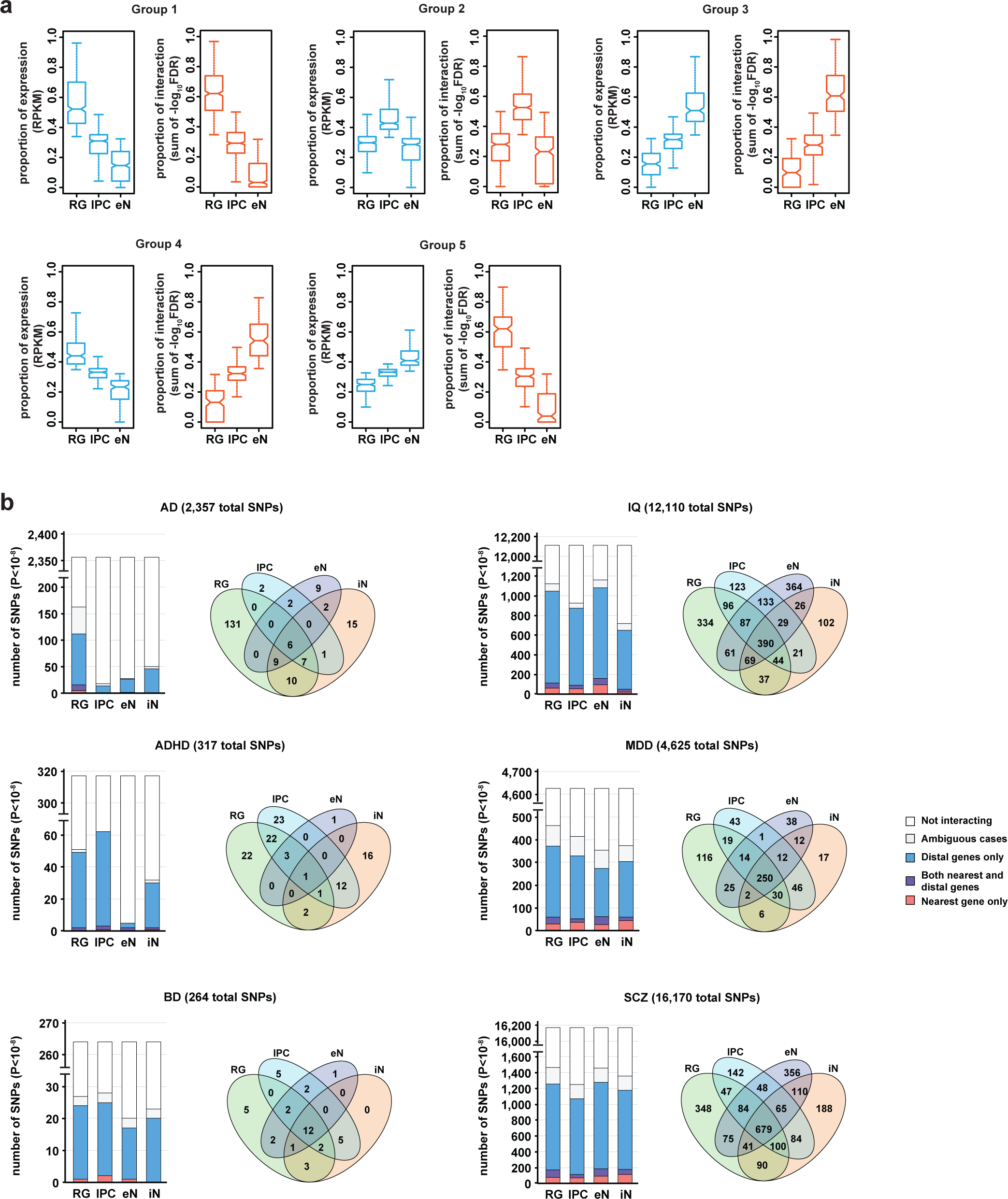
Developmental trajectories and annotations for complex neuropsychiatric disorder- and trait-associated variants. (**a**) Box plots showing the distributions of gene expression and cumulative interaction scores for groups in **Fig. 4**. The median, upper and lower quartiles, minimum, and maximum are indicated. (**b**) Bar graphs showing the numbers of GWAS SNPs (*P* < 10^-8^) interacting with their nearest gene only, with both their nearest and distal genes, or with distal genes only for each cell type and neuropsychiatric trait. Venn diagrams display the cell type-specificity of all interacting GWAS SNPs for each neuropsychiatric trait.

Supplementary Table 1. Sample metadata.

Supplementary Table 2. PLAC-seq, ATAC-seq, and RNA-seq data processing metrics.

Supplementary Table 3. Enriched GO terms for genes participating in cell type-specific interactions.

Supplementary Table 4. Motif enrichment at cell type-specific distal interacting regions.

Supplementary Table 5. Super interactive promoters for each cell type.

Supplementary Table 6. Enriched GO terms for genes associated with specific developmental trajectories.

Supplementary Table 7. Target gene annotation for enhancers overlapping chromatin interactions.

Supplementary Table 8. Target gene annotation for complex neuropsychiatric disorder- and trait-associated GWAS SNPs overlapping chromatin interactions.

Supplementary Table 9. sgRNA sequences used for functional validation.

